# Preoptic transcriptomes during courtship reveal the molecular basis of behavioral coordination in a highly social cichlid

**DOI:** 10.64898/2026.09.28.755186

**Authors:** Isaac Miller-Crews, Hans A. Hofmann

**Affiliations:** Department of Integrative Biology, The University of Texas at Austin, Austin, TX; Institute for Neuroscience, The University of Texas at Austin, Austin, TX

**Keywords:** RNA-seq, gene networks, Preoptic area (POA), social behavior, reproduction

## Abstract

Human and non-human animals often coordinate their behavior, a phenomenon thought to strengthen relationships by enabling sequential assessment of an interaction partner’s quality as a mate or rival, a process that directly affects evolutionary fitness. There is increasing evidence that behavioral coordination is reflected by synchronized neural activity of the interacting partners, though few studies have examined synchronization at the level of gene expression. Here, we use Burton’s Mouthbrooder cichlid, *Astatotilapia burtoni*, to examine how neural transcriptomes are synchronized while male-female pairs engaged in variable amounts of reproductive behavior. *A. burtoni* females naturally display highly coordinated reproductive behavior with a male partner only directly prior to spawning, yet this behavior can be experimentally induced with exogenous prostaglandin F2 alpha (PGF2α), which also acts by releasing pheromones to promote male courtship behavior. We quantified reproductive behavior and circulating sex steroid hormones of PGF2α-treated pairs. We then used transcriptome analysis of the preoptic area (POA) – a key node in the regulation of reproductive behavior across vertebrates – and identified gene co-expression modules that showed sex-specific activity in modulating reproductive behavior. Our results, which implement a novel approach to measure transcriptome synchronization, demonstrate a remarkable degree of male-female POA transcriptome synchronization that depends largely on the reproductive responsiveness of the female, mirroring how natural reproduction is driven by female behavior. Together, our work advances understanding of the molecular mechanisms by which interacting individuals align their actions and how they experience social interaction.

## INTRODUCTION

Sexual reproduction requires animals to be in the right place, at the right time, and in the right state. This coordination of behavior and physiology is regulated by brain regions with evolutionarily conserved expression of sex-steroid hormone receptors and neuropeptide hormones (O’Connell & Hofmann, 2011). As with all genes, these neuroendocrine pathways do not function in isolation and therefore are best understood within the broader context of the transcriptome to fully appreciate the suite of neural mechanisms that regulate reproduction. By combining reproductive behavior and transcriptomics, we can enhance our understanding of how individuals interact to successfully reproduce together.

Recent research has shed light on the degree to which simultaneous interbrain correlation, or synchronization, of neuronal activity occurs between socially interacting individuals through the coordination of their behavior (Coleman et al., 2021; Dumas et al., 2010; Gustison et al., 2024; Kingsbury et al., 2019; Liu et al., 2018; Zhang & Yartsev, 2019). Interpersonal synchrony across sociobiological systems is considered crucial in social relationships (Liu et al., 2018). Neural synchrony, as is observed in the mother-infant bond, is the result of multisensory signals and can reflect affiliation (Endevelt-Shapira et al., 2021; Reindl et al., 2018). In fact, *in vivo* calcium imaging simultaneously in the dorsomedial prefrontal cortex of socially interacting mice showed how neural synchronization of individual neurons results from continuous social behavior and is even predictive of future social interactions (Kingsbury et al., 2019). Likewise, the use of wireless electrophysiology in pairs of bats across various natural social interactions illustrated that the degree of neural synchronization covaries with the amount of social interaction and manifests across timescales, from minutes to hours (Zhang & Yartsev, 2019). Beyond neural activity, work in the Siamese fighting fish, *Betta splendens*, demonstrated that the individuals within a pair of fighting fish have higher neural transcriptomic synchronization when compared to fighting fish not within the pair (Vu, 2020). Importantly this transcriptomic research focused on males performing highly coordinated and ritualized aggressive displays and hypothesized that this synchronization is the result of a combination of aggression related gene pathways and capture of immediate-early gene cascades at the moment of collection. Whether this transcriptomic synchronization occurs in socially coordinating individuals experiencing and displaying different behavior, such as in sexual reproduction, remains an open question.

Social reproductive behavior necessitates behavioral coordination of mates, hence the social aspect, and is particularly important as this joint cooperation can impact fitness (Axelrod & Hamilton, 1981). During sexual reproduction, two individuals with different life histories function together while performing distinct actions. In response to reproduction, previous research demonstrates how activation patterns across neural networks overlap and differentiate across the sexes (Field et al., 2018; Field & Maruska, 2017). These neural networks occur in brain regions such as the preoptic area (POA), which is well known to play a major role in integrating external environmental cues with internal state to control social behavior (P. Chen & Hong, 2018; Dominguez & Hull, 2005). Even though the POA is activated during reproductive behavior in both males and females, this does not mean that the same underlying neuromolecular mechanisms are engaged, especially given the life history trajectories and current state of the animals (De Vries, 2004). Analysis of such mechanisms is made possible by RNA sequencing (RNAseq), which enables the discovery of the neural response at the level of gene expression. By applying RNAseq in both individuals while they are engaged in reproductive behavior, we can begin to understand what role transcriptomic synchronization might play in behavioral coordination between mates.

Burton’s Mouthbrooder, *Astatotilapia burtoni*, is a highly social cichlid fish and an ideal model system for studying neural response to reproductive opportunity. This is because both male and female *A. burtoni* demonstrate distinct reproductive states, as represented by commonly occurring correlated changes in behavior and physiology, which are regulated by and responsive to the social environment. As such, sexual reproduction in this species requires coordination of mates at both a physiological and behavioral level, to ensure that both partners are in the right place, at the right time, and in the right state. In *A. burtoni*, male-typical reproductive behavior is limited to dominant individuals, while females typically only ever interact with dominant males while gravid, i.e., carrying eggs. In turn, males exhibit increased amounts of reproductive behavior in direct response to reproductively active females (Huffman et al., 2012). As such, males display more courtship behavior (e.g. leads and displays) to entice reproductively ready females into their bower to mate, which is the only time females will leave the shoal to associate with dominant males. This stereotyped female reproductive behavior can be induced by treatment of females with a naturally occurring fatty acid hormone, Prostaglandin F2alpha (PGF2α), regardless of reproductive state (Kidd, Dijkstra, et al., 2013). We can therefore take advantage of the natural behavior of dominant males and hormonally treated females to reliably generate coordinated reproductive behavior in *A. burtoni*. While previous work in *A. burtoni* has demonstrated both shared and sex-specific responses across brain regions in response to reproduction, the POA is known to play an important role in both sexes (Field et al., 2018; Field & Maruska, 2017; Juntti et al., 2016; O’Connell et al., 2013). Yet, as the individual sexes are responding to and producing unique responses during reproduction, the mechanism driving these differences likely occurs at a yet untested biological level, such as in the transcriptome.

In the present study, we use 3’ Tag-based RNAseq to assess the transcriptomic response in the POA to reproductive opportunity between sexual partners of *A. burtoni*. Our study provides a framework to test if reproductive opportunity activates the same or different gene networks in a sex-specific manner. We hypothesized that as partners engaged in increased reproductive behavior and physiological coordination during a reproductive opportunity (‘High coordination’), so too would the degree of neural transcriptomic synchronization increase from these gene networks between males and females (Fig. 1). Specifically, this transcriptomic synchronization could occur due to relative similarity of partners in the activity of coordinated gene expression programs or across global transcriptomic state. Our work provides novel insight into how coordination at the behavioral and physiological level is integrated into synchronization at the neural transcriptomic level.

**Figure 1:**
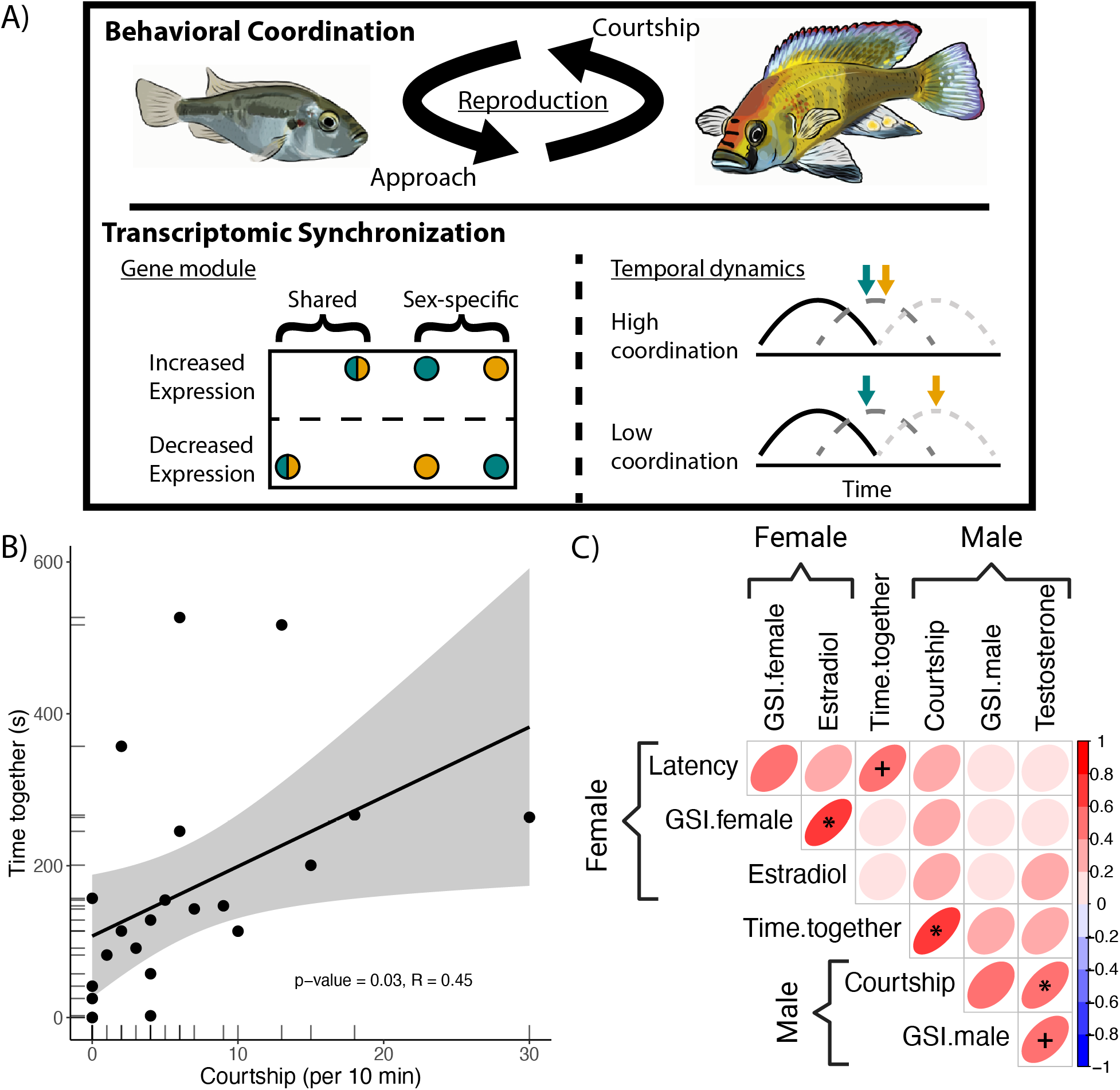
A) Graphical abstract of behavioral coordination and potential mechanisms of transcriptomic synchronization. <u>Top</u>: Reproduction requires both partners to engage and coordinate, only initiated through a combination of male courtship or female approach. Transcriptomic synchronization between partners could arise via coordinated expression of gene co-expression modules (<u>Bottom left</u>) or temporal dynamics of gene expression pathways (<u>Bottom right</u>). For gene co-expression modules, joint reproductive behavior between females (teal) and males (gold) could lead to correlated module response being either shared or sex-specific. For temporal dynamics, each reproductive pair will have differential coordination in the engagement of behavior. At the moment of collection, transcriptomic synchronization may arise due to timing of gene expression cascades triggered by the onset of the behavior occurring at the same time (‘High coordination’), while partners with less behavioral coordination may be at different points in the gene expression cascade due to differences in when each individual began engaging with their partner (‘Low coordination’). B) Male and female pairs demonstrate significant correlation of reproductive behavior as they coordinate during the reproductive assay. C) Correlation across all behavior and physiology traits of male and female partners during the reproductive assay (‘*’ = FDR adjusted p-value < 0.05, ‘+’ = FDR adjusted p-value < 0.1).

## METHODS

### Animal handling

Animals in this study came from a laboratory population descended from a wild-caught stock (Fernald & Hirata, 1977). All work was done in compliance with the Institutional Animal Care and Use Committee (IACUC) at The University of Texas at Austin.

### Reproductive behavior assay

All *A. burtoni* were housed in 120 L aquaria with communities of two males, dominant and subordinate, and three size-matched females separated by a clear divider from another community in the same aquarium. Social status of males was determined by coloration, behavior, presence of egg spots, and apparent blue lips. Experimental animals of both sexes were later confirmed via gonadal inspection. Communities were monitored via live global scoring between 10:00-12:00 every other day for a minimum of two weeks to ensure dominant males maintained a stable status and each community had at least one female that spawned and brooded successfully.

To test both sexes’ response to a reproductive opportunity in *A. burtoni,* we adapted a previously described reproductive behavior assay (Kidd, Dijkstra, et al., 2013; O’Connell et al., 2013). The experimental aquarium (100 L) was separated into three equal sized compartments with a clear barrier covered by an opaque divider that could be removed. The central compartment contained one terracotta pot that served as territorial shelter for the dominant male while the female was placed in a lateral compartment. At the front of the aquarium, a joint bower was placed across each barrier to serve as a focal location during the behavioral trial. In the afternoon prior to the experiment between 16:00-18:00, a dominant male and a female, who was selected for not being visibly gravid or brooding and within 1 cm standard length of the male, from separate aquaria were acclimated overnight in the experimental aquarium. On the morning of the experiment, after the overnight aquarium acclimation, all fish were fed at least one hour prior to the experiment. Each female was then injected with a biologically relevant dose of PGF2α to induce reproductive behavior (Kidd, Dijkstra, et al., 2013), returned to its respective experimental aquarium, the black opaque dividers removed, and video recorded for one-hour.

After a five-minute recovery period from the injection, reproductive behavior of both sexes was scored from video using BORIS during a 10-minute window immediately following when the female first engaged with the male. The main behaviors scored were latency for the female to engage (latency), male courtship directed at the female including quivering and lateral displays (courtship), and how much time the females and males spent together at the bower (time together).

### Tissue processing

Immediately following the one-hour behavior trial, animals were euthanized by rapid cervical decapitation. Body mass, gonad mass, and body length were measured, and brain, gonads, and blood were harvested. Gonadosomatic-index (GSI), calculated as the ratio of gonad mass divided by total animal mass (Giese, 1959), was used as a proxy for reproductive condition. Extraction of plasma from blood occurred shortly after dissection before being frozen at −80 C for later steroid hormone analysis. Circulating sex steroid hormones, estradiol for females and testosterone for males, were assayed by ELISA (Cayman Chemicals, USA). Brains were flash frozen in OCT and stored at −80 C prior to being cryo-sectioned at 300 µm for later micro-dissection. The POA was micro-dissected following standard procedures established in the Hofmann lab. RNA was extracted with automated Maxwell RSC simplyRNA tissue kit (Promega, USA).

### Transcriptome sequencing

The development of cost-effective, and yet highly accurate, RNAseq methods such as the 3’ Tag-based approach expands the opportunity to study transcriptomics across a broader range of contexts than prior more expensive approaches (Lohman et al., 2016; Meyer et al., 2011). After quality control on the Bioanalyzer, all 44 samples were submitted to GSAF for library preparation and 3’ Tag-based RNA sequencing (Meyer et al., 2011). Read quality was assessed with FastQC (Andrews, 2010) and MultiQC (Ewels et al., 2016). The transcriptomic data was referenced to the closest relative of *A. burtoni* with the highest-quality genome, the Nile Tilapia (O_niloticus_UMD1) (Conte et al., 2017), using the STAR aligner for mapping and counting (Dobin et al., 2013). All forty-four samples passed QC filters for analysis for a total of 22 male and female pairs. We obtained on average 4.1 million reads per sample and had mapping efficiency of ∼82.63% uniquely mapped aligned reads to the Nile Tilapia genome. Genes were filtered to a minimum read count of at least two and present in half the samples. A total of 13,343 genes were represented following threshold filtering (Dataset S3).

### Statistical analysis

Behavioral and physiological data for each pair were analyzed for correlations and assessed with principal component analysis (PCA). Correlation analysis was done with rcorr (R packages: Hmisc (Harrell Jr. & Dupont, 2019)) and corrected for false discovery (FDR) with rcorr.adjust (R packages: RcmdrMisc (Fox et al., 2023)).

We identified differentially expressed genes (DEGs) between males and females using DESeq2 to normalize and quantify gene expression with a FDR p-value <0.1 (Love et al., 2014). We used PCA of the top 1000 most variable genes to compare the distribution of males and females.

Weighted gene co-expression network analyses (WGCNA) with all samples allowed for unbiased identification of modules of genes with correlated expression (Langfelder & Horvath, 2008). WGCNA with both sexes together was used to identify the presence of any sex specific modules or modules responsive to reproductive opportunity in either sex. First, we tested for sex specific modules by calculating and comparing the 95% confidence interval of module eigengene (MEG) values for each sex in a forest plot. Next, we used FDR corrected spearman rank correlation on MEG values with sex-specific behavior and physiological traits, as well as between partners, to identify modules that showed correlated expression. Gene module membership was compared to a list of known immediate-early genes and list of candidate genes related to social behavior (Dataset S5). Putative functional roles of WGCNA modules were assigned with Gene Ontology (GO) term enrichment. We used AnnotationForge to generate a GO database for Nile Tilapia (Carlson, M & Pagès, H, 2024), calculated GO enrichment with enrichGO from the package clusterprofiler (Wu et al., 2021), with FDR corrected adjusted p-value cutoff of 0.05, minGSSize = 10, and maxGSSize = 500, and graphed our results the package enrichplot (Yu G, 2025).

To test for transcriptomic synchronization, we developed an analysis approach comparing transcriptomic similarity between an individual and their partner to all other opposite-sex partners. We used two approaches to calculate transcriptomic similarity: a PCA approach and a MEG-based approach. First, for the PCA approach, we performed PCA of the normalized gene expression data with all samples. Then, for each individual we took the PC scores of the top PCs that accounted for up to 80% cumulative variance. We calculated transcriptomic distance between all individuals as the Euclidean distance of PC scores weighted by the variance of a given PC, therefore giving greater weight to distance on PCs that explained more variance of the data. This PCA weighted-Euclidean distance gives a transcriptomic similarity score that reflects global transcriptomic state. For the second transcriptomic similarity measure, we used MEG scores from WGCNA and calculated distance between individuals. This MEG distance gives a transcriptomic similarity score that reflects shared co-expression program activity. Using both the PCA- and MEG-based distance measures separately provided a distribution for each individual to every possible opposite-sex partner, including the actual partner. We then calculated a z-score across each individual for their respective partner to calculate a transcriptomic synchronization score, with higher values indicating that an individual was more similar to its partner relative to any random individual of the opposite sex.

Next, for both the PCA- and MEG-based transcriptomic synchronization scores, we used a decision tree method to enable the identification of which variables were most relevant to determining an experimental outcome (White et al., 2023), in this case, the degree of transcriptomic synchronization. These methods are useful for imputation in that they are robust to outliers, work on a range of variable types, handle intercorrelated predictors, function even when used with relatively small n and large p (e.g. limited sample size and large number of variables), and generally handle cases when assumptions for regression are not met (Levshina, 2020). We applied this approach to determine how different variables of a given individual affected the degree of transcriptomic synchronization for both the PCA- and MEG-based scores. Specifically, we used Conditional Inference Trees (CITs) to determine the effect of a predictor variable on transcriptomic synchronization while simultaneously considering the effects of all other predictor variables (Hothorn et al., 2006). Briefly, CITs select the predictor that best distinguishes differences in the response variable before using p-values, with a Bonferroni correction, to determine if the predictor is significant, in which case the variable is split, and the data is partitioned. This occurs recursively until the pre-defined minimum split size or node size is reached (Hothorn et al., 2006). We assessed the impact that the behavior and physiology traits of both the individual and partner have on transcriptomic synchronization score for each sex independently using the R packages party and partykit (Hothorn et al., 2023, 2026). A significance threshold of 0.1, minimum splits ten, and minimum node size five was used for tree construction. For both the male and female models, and for both the PCA- and MEG-based transcriptomic synchronization scores, the following variables were used: time together, latency, female GSI, male GSI, female estradiol, male testosterone, and courtship. Variable importance between the sex-specific and within sex models across the two transcriptomic synchronization scores were compared to test how different predictors affected the level of synchronization both between each sex and between the two measures of transcriptomic synchronization within a sex.

## RESULTS

### Behavior and physiology covary between males and females during reproductive behavior

We used a paradigm of induced reproductive behavior in pairs of *A. burtoni* to measure behavioral and physiological coordination between partners. Of all the pairs, 20/22 engaged in some amount of affiliative behavior or social interaction. We predicted that there would be a correlation across male-typical reproduction associated traits (courtship, male GSI, and male testosterone) and across female-typical reproduction associated traits (time together, latency, female GSI, and female estradiol), as these occurred within the same animal. Importantly, our behavioral assay resulted in a significant correlation between male courtship behavior and time together (R^2^ = 0.2043, F(1, 20) = 47.99, p =.0.03) (Fig. 1b). As predicted, in comparisons across all behavior and physiology traits for a pair, we found a significant Spearman’s rank positive correlation between female GSI and female estradiol (r_22_ = 0.5893, p_adj_ = 0.039) and a positive albeit non-significant correlation between male testosterone and male courtship behavior (r_22_ = 0.6453, p_adj_ = 0.0741) (Fig. 1c; Dataset S2).

### Differential gene expression analysis reveals only small differences between sexes

To determine the degree of sex differences present in the POA transcriptome, we used differential gene expression analysis to determine the number of sexual biased genes. We identified only 13 differentially expressed genes (DEGs) across sexes at an FDR-adjusted p<0.1, with the strongest female-biased gene being *agouti-related protein* (*agrp*) and the male-biased gene being *oxytocin* (*oxt*) (Fig. 2a). Additionally, PCA did not reveal any sex differences for any of the PCs (Fig. 2b).

**Figure 2:**
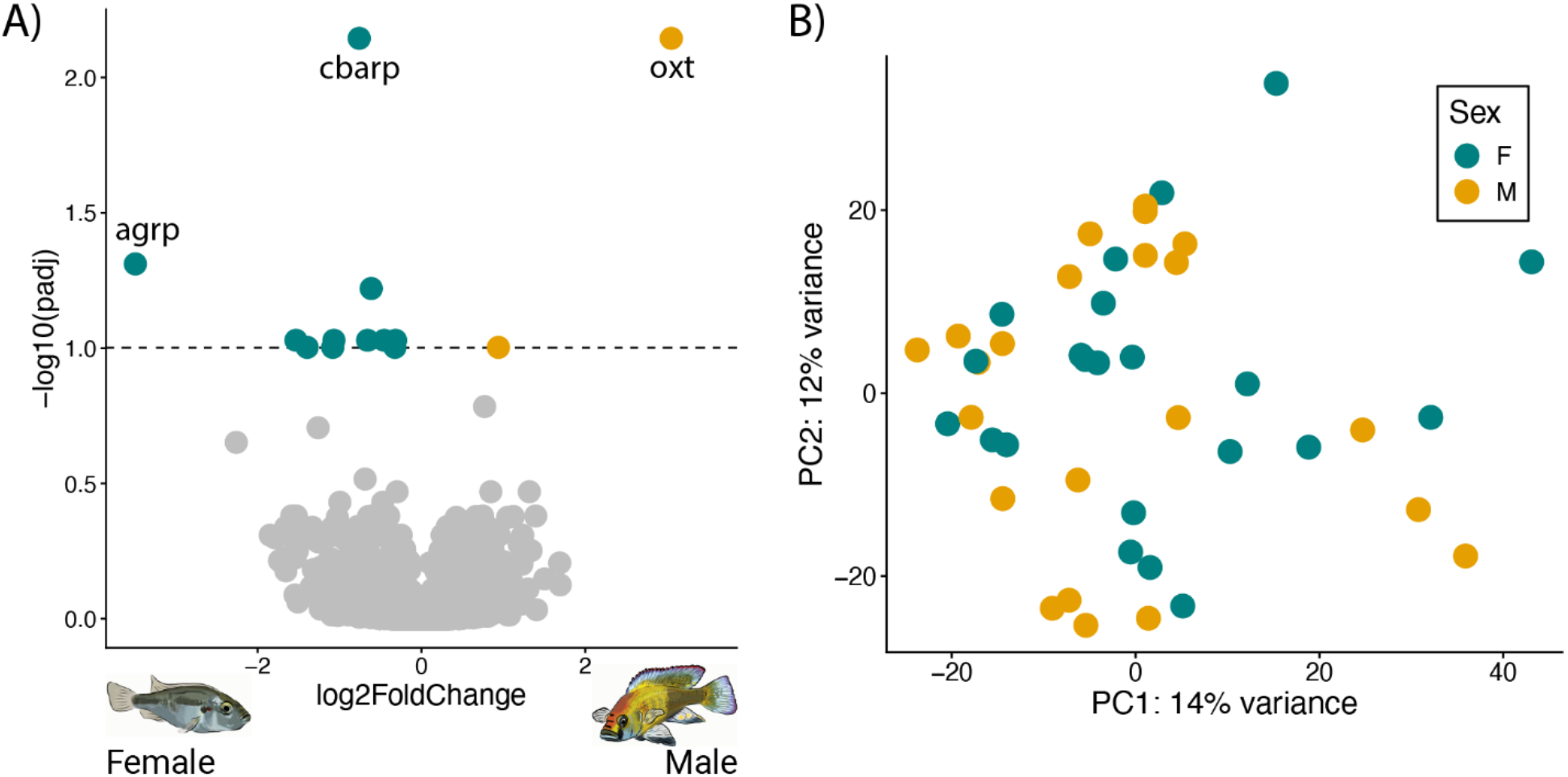
A) Differential POA gene expression between males (gold) and females (teal) following reproductive assay. B) Males and females show no clear separation in PCA distribution for POA gene expression following reproductive assay.

### Transcriptomic principal components show weak correlations with behavior and physiology across sexes

When we next compared the transcriptomic PC scores with the behavior and physiology data, we found no significant correlations that passed FDR < 0.1 between PC1, PC2, and PC3 (which account for 14%, 12%, and 10% of variance, respectively), indicating a wide range of natural variation (Fig. S2a). Before FDR correction, we did find significant positive correlations between PC1 score in females and both estradiol level and time spent together, while in males we did find a significant positive correlation between PC1 and courtship, PC3 and male GSI, and a negative correlation between PC2 and time spent together. We further tested whether the PCs reflected interbrain correlation between partners and found no significant correlations of the PC scores for PC1 or PC2, but we did find a positive albeit non-significant correlation of PC3 (r_22_ = 0.36, p = 0.077) (Fig. S1b). While none of these correlations passed FDR adjustment, these results point toward the possibility of a more complex relationship between phenotype and transcriptomic state as measured via PCA.

### Gene co-expression networks

Genes do not function in isolation, rather within the context of the entire transcriptome. We can harness the nature of these gene co-expression networks to identify potentially functional modules of genes while reducing noise and transcriptomic dimensionality. WGCNA across all samples together allowed for the identification of potentially sex-biased modules or gene networks responsive to reproductive opportunity in each sex. We successfully identified 10 gene co-expression modules in the POA of males and females (Fig. 3a; Fig. S2). We found no evidence for sex-specific gene co-expression modules (Fig. S3a). We tested MEG expression in males with male traits, and only the magenta module displayed a positive yet non-significant correlation with courtship behavior (r_22_ = 0.408, p_adj_ = 0.059) (Fig. 3c). Similarly, in females, when comparing correlation with female traits, time spent together was positively but not significantly correlated with both the blue module (r_22_ = 0.611, p_adj_ = 0.076) and the purple module (r_22_ = −0.59, p_adj_ = 0.076) (Fig. 3d).

**Figure 3:**
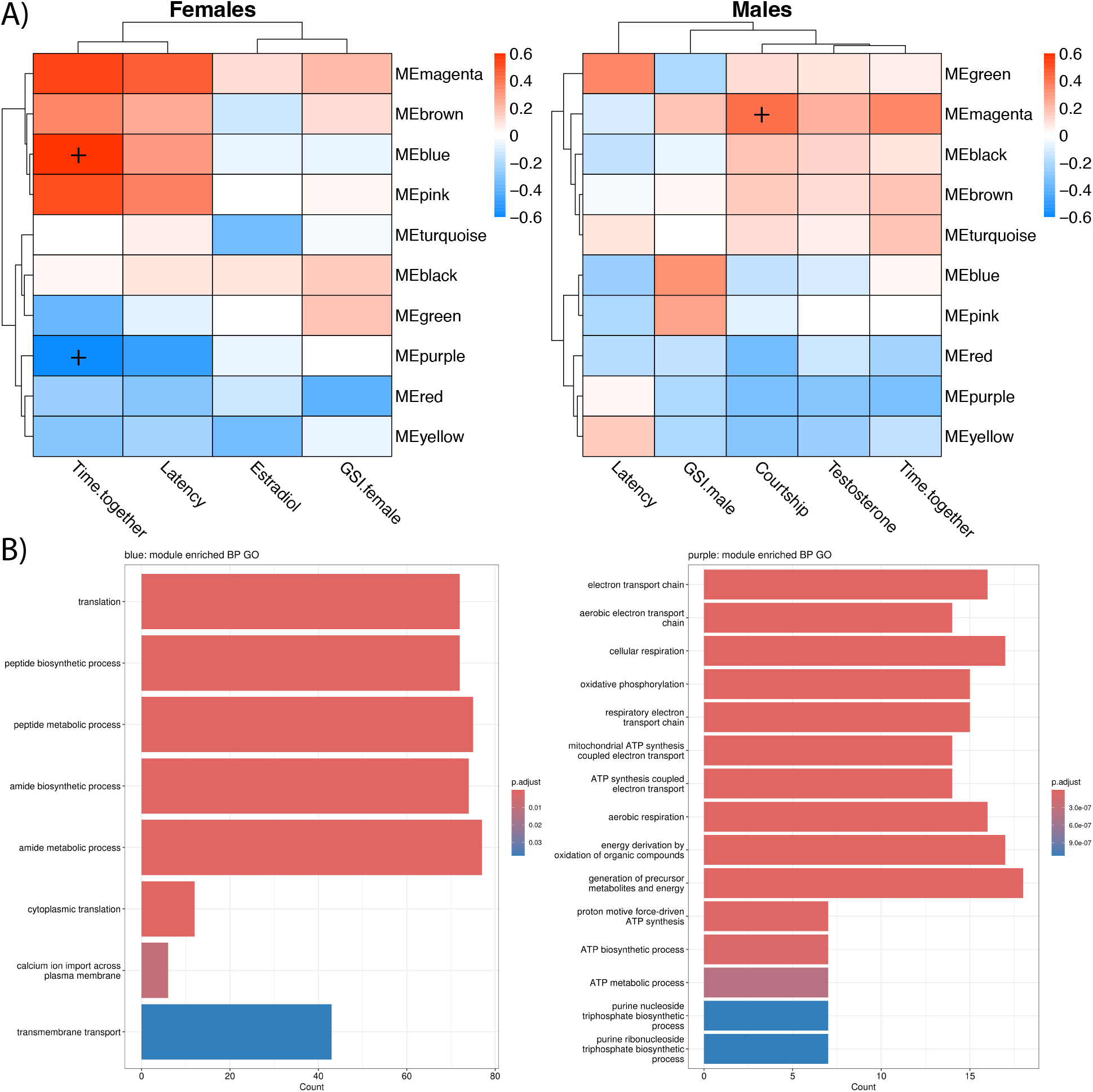
A) Correlation heatmap of sex-specific module eigengene expression with sex-specific behavior and physiology traits (‘+’ = FDR adjusted p-value < 0.1) in females (left) and males (right). B) GO term enrichment bar plot for significant modules blue (left) and purple (right), with number of enriched gene (x-axis) and FDR adjusted p-value (color). There are no significantly enriched GO terms for magenta module.

We used GO term enrichment analysis to assign putative functional roles to WGCNA modules using gene module membership (Fig. 3b; Fig. S3b; Dataset S4). We found no significant enrichment for either the green or magenta modules. We found significant enrichment of biological processes with top GO terms in the pink module for immune system process and immune response; in the black module for cell population proliferation, microtubule-based process, cilium movement and axoneme assembly; in the blue module for translation, transmembrane transport, and calcium ion import across plasma membrane; in the purple module for oxidative phosphorylation and aerobic electron transport chain; in the yellow module for multicellular organism development, cell differentiation, hormone transport and secretion; in the turquoise module for protein-containing complex organization and mRNA processing; in the brown module for regulation of molecular function, neural crest cell migration, and mesenchymal cell migration; and in the red module for response to growth factor, brain development, and cellular response to growth factor.

We hypothesized that modules associated with reproductive behavior contained immediate-early genes (IEGs), indicating neural activation within the POA in response to the reproductive opportunity, or candidate genes identified in previous research as associated with social behavior, including reproduction. Of the 12 IEGs tested, half were present in at least one module, with three being in the turquoise module, and only one gene associated with an apparent reproduction associated module, *klf8*, in the purple module (Dataset S5). Similarly, for the candidate socially related genes, six (*htr2cl1*, *nr3c2*, *hsd11b2*, *htr1aa*, *HTR1F*, *gnrh3*) were contained in the blue module, which was positively associated with female reproductive behavior, and three (*glula*, *OPRK1*, *oprm1*) were contained in the magenta module, which was positively associated with male reproductive behavior. Of the 50 candidate genes, 16 genes (*agrp*, *ara*, *avp*, *crhbp*, *DLK1*, *gnrh1*, *POMC*, *CRH-BP*, *esr1*, *esr2a*, *gal*, *NR3C1*, *oxt*, *pdyn*, *trh*) were in the yellow module (Dataset S5).

### Transcriptomic synchronization

One way in which individuals could synchronize transcriptomic activity is through coordinated co-expression patterns, with either activation of shared modules or correlation of sex-specific module activation (Fig. 1b). Although not significant after FDR correction, we found evidence that the black module may represent a shared activation module between partners (Fig. 4). Likewise, several modules showed a trend towards correlation between partners, with the strongest effect found in the negative correlation between MEG expression of the red module in males and black module in females (Fig. 4).

**Figure 4:**
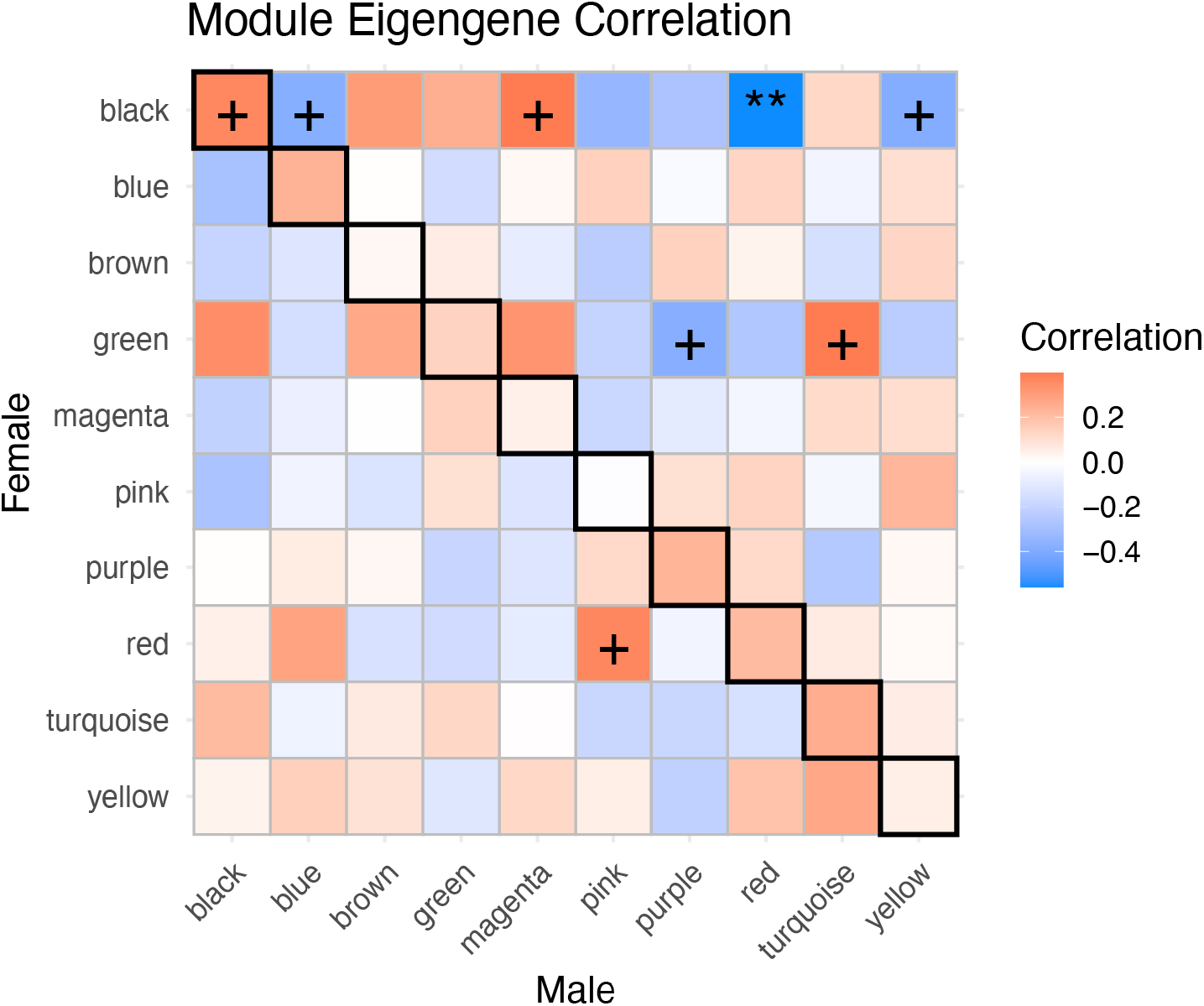
Correlation heatmap of module eigengene expression across pairs of males (x-axis) and females (y-axis) (‘+’ = p-value < 0.1, ‘*’ = p-value < 0.05, ‘**’ = p-value < 0.01). Dark border indicates correlation of same module between partners.

We hypothesized that the level of behavioral coordination any pair exhibited would be reflected in the degree of their POA transcriptomic synchronization (Fig. 1b). To test this, we developed two methods for assigning a pairwise similarity score as a metric for transcriptomic synchrony (see methods). The first method uses weighted-Euclidean distance of gene expression PCA to measure similarity in global transcriptomic state between individuals. The second method uses distance of MEG expression to measure similarity in coordinated gene expression patterns between individuals. Both the PCA- and MEG-based distance scores were then converted to a transcriptomic synchronization score as the z-score between an individual and their partner compared to every other opposite-sex individual. Though the similarity score for a male and female from the same pair are related, the values differ as the male similarity score to their female partner is relative to the distribution of all females, and visa versa for females to males. As the distribution of males and females differ, this results in the similarity score for a male and female from the same pair being slightly positively correlated, but not significantly (Fig. S5). Unsurprisingly, we found a significant correlation between the PCA and MEG transcriptomic similarity scores of partners (r_22_ = 0.92, p < 0.001) (Fig. S5). We found for both approaches that pairs of individuals displayed a range of relative transcriptomic similarity scores across partners compared to all other individuals of the opposite sex (Fig. S4a, S4b). We, therefore, did not find higher transcriptomic similarity between an individual and every other opposite sex-individual, for either males or females (Fig. S4c, S4d). Since individual pairs vary in behavior and physiology, this lack of general similarity indicates that shared environment alone is not enough to drive transcriptomic synchronization.

The complex interaction between all the various behavior and physiological traits ultimately produces the individual phenotype and response. We therefore attempted to test the degree to which phenotypic traits predicted transcriptomic similarity for each sex with Conditional Inference Trees (CIT). CIT analysis was performed to recursively determine the significance (Bonferroni corrected p-value < 0.05) and effect of a predictor variable while simultaneously considering all other effects. For each sex, we were able to generate a tree and predictive model of the behavior and physiology traits on the degree of transcriptomic synchronization for both the PCA- and MEG-based scores (Fig. 5). For the PCA-based transcriptomic synchronization score, in females we found a significant effect of female GSI, followed by male courtship, while in males we found a significant effect of female estradiol level, followed by female gonad size, and then male courtship (Fig. 5a; Fig. S6a). For the MEG-based transcriptomic synchronization score, in females we found a significant effect of male courtship, followed by male testosterone, and then time spent together, while in males we found a significant effect of female estradiol, followed by latency for female to engage with male (Fig. 5b; Fig. S6b).

**Figure 5:**
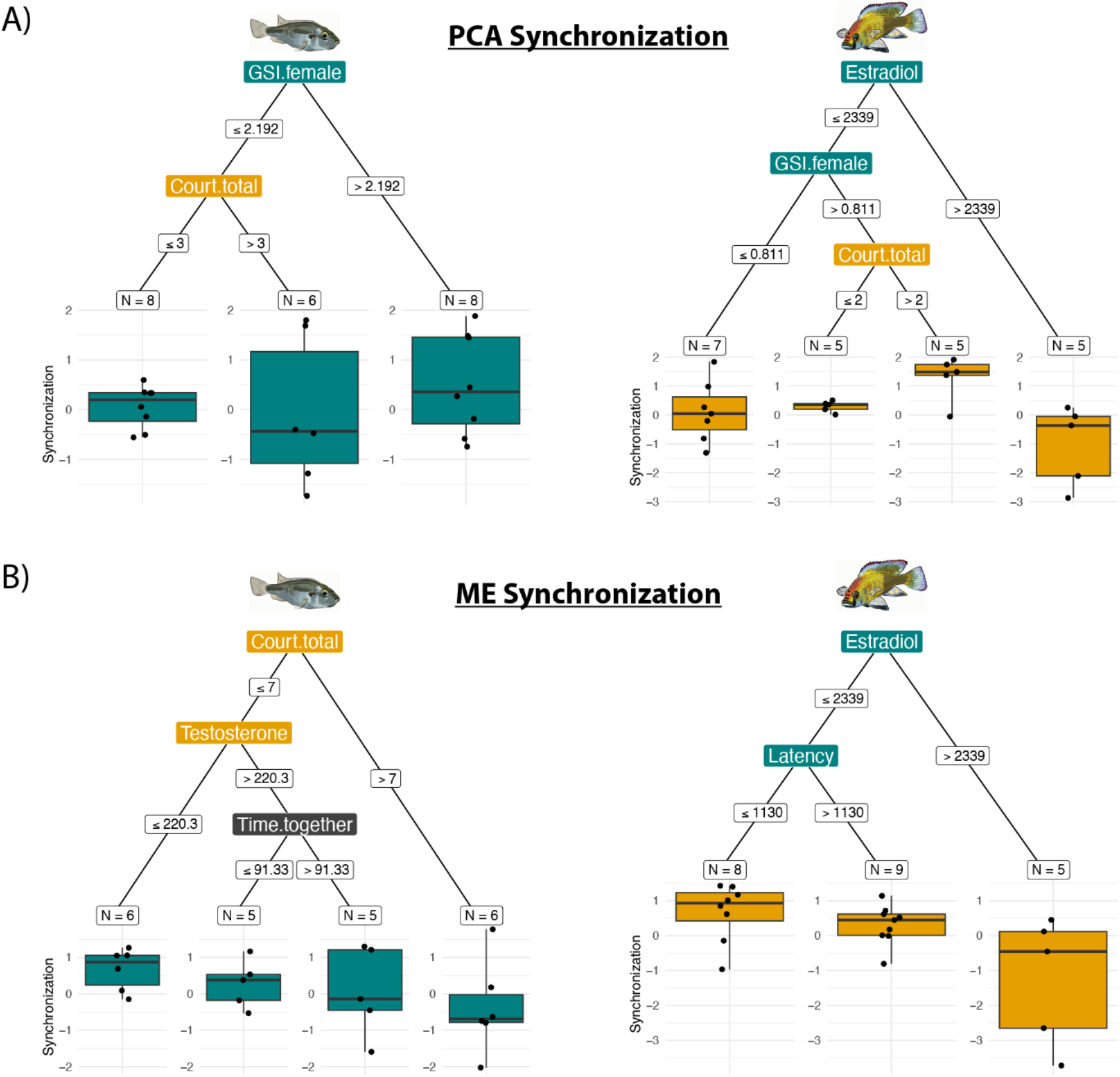
Conditional inference trees for predictors of transcriptomic similarity as measured through A) weighted-Euclidean PCA distance of gene expression or B) WGCNA module eigengene distance. For both the PCA and MEG distance measures, the degree of synchronization (boxplots) was assigned for an individual as the z-score of distance measures between a female and their male partner compared to every other male (left, teal boxplots), or for a male relative to every other female (right, gold boxplots). A higher synchronization score indicates more similarity to their partner relative to all other opposite sex individuals. Starting from the most impactful variable at the top through each success significant variable (alpha < 0.1), each node in the decision tree represents a significant split among samples with the node labeled by the predictor and the edges labeled as the split value. Node labels are colored by whether they are attributed to the male (gold), female (teal), or combination (grey).

The variable importance of the male- and female-specific models was compared across the PCA and MEG synchronization score models to identify which variables have a larger sex-specific impact on predicting transcriptomic synchronization. For females, the largest differences in relative variable importance between the models indicates that female gonad size is important for relative PCA similarity, while male courtship and testosterone are important for relative MEG similarity (Fig. S6C). For males, both the PCA and MEG synchronization scores had female estradiol as the most important variable.

## DISCUSSION

In the present study, we demonstrated how the POA transcriptome of *A. burtoni* reflects response to reproductive opportunity in both sexes and the degree of transcriptomic synchronization that arises from the resulting coordinated behavior (Fig. 1). To achieve this, we implemented a reproductive behavior assay that successfully correlated male and female reproductive behavior. Even without any major sex differences in the POA transcriptome, we discovered gene co-expression networks that regulate reproduction in a sex-specific manner. While we did not find evidence for global transcriptomic synchronization between partners, how much an animal is engaging in reproductive behavior is predictive of the degree of transcriptomic synchrony with its partner.

The POA is notable for its evolutionarily conserved functional role across vertebrate taxa as a key neural node in social behavior and as a regulator of reproduction (P. Chen & Hong, 2018; O’Connell & Hofmann, 2011). In rodents and humans, the POA is morphologically differentiated between the sexes, to the point that a subregion is designated as the sexually dimorphic nucleus of the preoptic area (SDN-POA) (Hofman & Swaab, 1989; Tsukahara & Morishita, 2020). Yet, most of this research has been focused on cellular characteristics and done in mammalian species, with relatively simplistic sex determination and sexual differentiation systems. Cichlid fish in general, *A. burtoni* included, exhibit a complex and apparent evolutionarily flexible set of sexual differentiation mechanisms (Behrens et al., 2022; El Taher et al., 2021; Roberts et al., 2016). Even in clades with highly conserved sex chromosomes, such as songbirds, sex differences in neural transcriptomes vary a great deal, with a majority of the differences emanating from the sex chromosomes (Miller-Crews et al., 2026). Among fish species, sex differences from whole-brain transcriptomes can ranges from the hundreds in species such as zebrafish (Wong et al., 2014) or tropical gar (Cribbin et al., 2017), to the thousands in species such as blue gourami (Degani & Meerson, 2024). While in *A.* burtoni, whole-brain analysis using microarray identified 316 male-biased and 253-female-biased genes (Renn et al., 2008). So, surprisingly, we did not find major sex differences in the POA transcriptomes of *A. burtoni* (Fig. 2). Additionally, our experimental design enhanced the natural variation by including a range of individuals from each sex. Yet, even with all this variation, only 13 genes were found to be sex-biased, with the top male-biased gene being *oxt* and the top female-biased gene being *agrp*. While research into *oxt* in *A. burtoni* is limited, within teleost fish there does appear to be a functional sex-specific role in relation to reproduction (Huffman et al., 2012; Mennigen et al., 2022), specifically with increased cellular expression in reproductively active males (Miller-Crews & Hofmann, 2026). Similarly, research into sex-differences of *agrp* gene expression is minimal, and in the case of chickens appears to be inverse to the effect found in this study (Caughey et al., 2018; Loganathan & Belsham, 2016).

Even in the absence of major sex difference, males and females have distinct gene co-expression modules that correlate with their respective reproductive behavior (Fig. 3a). This implies that sexual behaviors that are sex-typical are regulated by independent gene networks. Yet, the lack of sex difference for any of the gene modules demonstrates that they are not exclusive to either sex (Fig. S3a). Importantly, this is all occurring within the POA, which has been shown to be responsive to reproductive opportunity across sexes in *A. burtoni* (Burmeister et al., 2005; Field et al., 2018; Field & Maruska, 2017; Juntti et al., 2016; O’Connell et al., 2013), which is itself a highly conserved trait of the POA across vertebrates. Therefore, while this region is important for both sexes in regulating reproduction, how this manifests at the transcriptomic level appears to be a much more complex interaction. Work in mice has demonstrated that mating induces differential activation of specific transcriptomic cell types in the POA (Moffitt et al., 2018), while research in the POA of closely-related deer mice species has demonstrated how sex differences in cellular abundance may be conserved for specific transcriptomic cell types (J. Chen et al., 2025). Therefore, the mechanistic response of the POA to activation in a reproductive context is dependent on the role of the individual in the encounter.

Assessment of which genes and their associated GO terms are present in these gene networks associated with sex-typical reproduction provides insight into the nature of this mechanistic response. For example, the magenta module in males was most strongly correlated with male courtship behavior (Fig. 3) yet had no significantly enriched GO terms. While in females, two modules had opposite correlations with the amount of time a female spent with a mate (Fig. 3). This associative behavior in *A. burtoni* between dominant males and females only occurs in a mating context and is driven by female choice (Kidd, O’Connell, et al., 2013). For the blue module, which in females was positively correlated with association time, GO term enrichment highlighted regulation of both transcription and translation, hinting at the future engagement of gene pathways not captured with the temporal resolution of this experiment. For the purple module, which was negatively correlated in females with time spent together, GO term enrichment highlighted cellular respiration, which may indicate differential metabolic activity when females are more fully engaging in reproductive behavior.

To gain insight into the gene pathways engaged, we used a candidate gene approach. Surprisingly, the blue module contained *gnrh3*, which is known to be involved in reproduction, but the peptide is exclusively localized in the terminal nerve ganglion in fish (Grone et al., 2010; Hu et al., 2016). While it is possible that the microdissection punches for the POA captured a region more rostral than anticipated, we caution that gene expression is separate from mature peptide and it is likely that the gene shares transcription factors with other genes in the blue module. The blue module has multiple genes for serotonin receptors which have previously been shown to modulate social behavior in fish, including *A. burtoni* (Loveland et al., 2014; Stettler et al., 2021). Additionally, this module contained an enzyme and receptor involved in the glucocorticoid system, which has been implicated in relation to reproduction (Leary & Baugh, 2020), including work in treefrogs demonstrating a role of glucocorticoids in regulating female mate choice (Baugh et al., 2021). Together, this implies that correlated gene expression surrounding glucocorticoid signaling in *A. burtoni* also regulates female reproductive willingness to engage in reproductive behavior.

One module that stood out was the yellow module, as it contained a suite of “usual suspects” for genes known to be related with reproduction (Dataset S5). Surprisingly, this module was not associated with any behavior or physiological trait that we measured. One explanation for this result could be an indication of the amount of natural variation resulting from our experimental design. It is possible that since all individuals were reproductively mature adults and interacting socially to some degree, even if just for a short amount of time, this gene network would not be differentiated across our samples. Alternatively, reproduction functions across a series of timescales. Our experimental design was aimed at targeting the initial response to a reproduction opportunity and thus would not capture longer-term changes.

For all these sex-typical responses to reproduction, the two individual partners are engaging in joint mating behavior. In our study, we did not find evidence for wide-scale synchronization of transcriptomic state or gene network expression between reproductive partners, indicating that if this were occurring it is not just due to shared environment alone (Fig. S4). As has been previously shown, individuals engaging in social behavior together experience an increased degree of simultaneous interbrain correlation in neuronal activity synchronization (Kingsbury et al., 2019; Zhang & Yartsev, 2019). This will occur in humans even when those individuals are at drastically different developmental stages and are participating in unique ways (Dumas et al., 2010; Endevelt-Shapira et al., 2021). Likewise, research in prairie voles has shown that the amount of reproductive behavior correlates directly with the degree of neural activity synchronization during pair bond formation (Gustison et al., 2024). As far as transcriptomic synchronization goes, research in fish has demonstrated how two males engaging in reciprocal aggressive behavior will have this synchronization manifest at the level of the transcriptome (Vu, 2020). Therefoe, due to the role of social behavior in driving synchronization at other neurobiological levels, the lack of general transcriptomic synchronization between pairs in our study could be a consequence of the two partners being different sexes and performing different behavior during reproduction, resulting in differential mechanisms in the POA regulating sex-typical behavior independently.

One hypothesis we had was that partners may demonstrate correlated expression of specific modules across partners, representing a shared pattern if the correlation occurred for the same module in both sexes or a sex-specific pattern if the correlation was between different modules in each sex (Fig. 1b). We found evidence for both instances, although our sample size makes FDR correction prohibitive for the 100 comparisons across all pairs of 10 modules between sexes (Fig. 4). We identified that expression of the black module in females appeared correlated to multiple other modules in males, including a shared pattern with expression of the black module in males. While the black module displays no obvious relationship with any of the phenotypic traits in females, nor is the GO enrichment for cell population proliferation particularly telling, its relationship with expression of modules in another individual warrants future investigation into what these ∼300 genes could represent in other social contexts.

Our dual measures for transcriptomic synchronization span both global transcriptomic state and co-expression program activity, meaning that if transcriptomic synchronization is occurring, it is in response to a complex interaction of multiple variables. This necessitates trying to understand which variables do predict the degree of synchrony for each sex separately, with the use of a tool such as CIT. The generation of these CIT models implies that, to some degree, the phenotype of both the individual and partner can predict synchrony due to the recursive significance required.

What may be driving the degree of transcriptomic synchronization of the female appears to change when considering global transcriptomic state or activity of coordinated gene pathways (Fig. 5). For females, relative similarity of global transcriptomic state does not have an obvious predictor, while relative similarity of gene co-expression program activity does appear to be predicted by less reproductive behavior and lower sex steroid levels from the male. This implies that females have a higher degree of relative similarity that are not actively engaging in reproductive behavior, as both the male and female are likely performing the same behavior, e.g. neither fish is socializing and are stationary in the tank. This mirrors the previous results showing transcriptomic synchronization relates to individuals performing the same behavior (Vu, 2020), although here we demonstrate that this remains true even if the life history context of the individuals diverge (i.e. same vs opposite-sex individuals).

Unlike in the female, in which partner behavior was a main predictor, we found that males are relatively less similar to their female partner if their female partner has high levels of estradiol (Fig. 5). For males, relative similarity of global transcriptomic state is predicted by the female having lower levels of estradiol while still maintaining a high degree of reproductive potential in conjunction with higher male courtship behavior. Similarly, if a female does not have very high levels of estradiol and engages with the male in the reproductive assay more quickly, the male shares a relatively more similar co-expression activity program. This implies that how similar a male is to their partner is driven by the female’s reproductive state and motivation to engage with the male in the reproduction assay.

Together, when we look at the variable importance between the models (Fig. S6), we find that the degree of transcriptomic synchronization appears to be driven by the partner’s phenotype. What appears to be most important is how the female responds to the reproductive opportunity, as this is the rate-limiting step for both males and females. In our design, the degree to which the partners can coordinate behavior at all is determined by how responsive the female is towards the reproductive opportunity, which in turn is dependent on her reproductive state. In contrast, dominant male *A. burtoni* maintain a constant level of vigilance to reproductive opportunities and are, therefore, already primed for reproduction. As this mirrors natural reproductive behavior, this implies that it is variation among females in reproductive responsiveness that ultimately drives how the male neural mechanisms respond. This result underlies that sex-specific mechanisms are involved, although they are not necessarily limited to either sex, but rather that what type of reproductive behavior an individual is performing impacts neural response. This means that the degree to which an individual engages in behavioral coordination is likely reflected in the neural transcriptomic synchrony.

## CONCLUSIONS

Sexual reproduction is a social interplay between two individuals coordinating their behavior to ultimately produce offspring. In this study, we tested how the POA transcriptome of both sexes in *A. burtoni* responds to a mating context. Even though the same brain region is engaged in regulating this behavior across sex, we show that the underlying neuromolecular pathways reflect sex-typical behavior. The extent to which the neural transcriptome synchronizes between partners during mating appears to be, at least partially, dependent on the reproductive responsiveness of an individual in the social interaction. As reproduction is the result of each individual partner producing and receiving different cues, e.g. only the male does courtship and receives female physiological cues, whether this maintains across other social contexts remains to be seen.

## DATA AVAILABILITY STATEMENT

The filtered reads have been deposited to the NCBI Short Read Archive (SRA), bioproject XXXXXX. The code used in this paper together with documentation can be accessed as the GitHub repository, https://github.com/imillercrews/Burtoni_repro_opportunity.

## ACKNOWLEDGEMENTS

We thank members of the Hofmann Lab for guidance and discussion. We also thank Dr. Liz A. Aguilar and Dr. Tessa K. Solomon-Lane for helpful suggestions and feedback on an earlier version of the manuscript. Sequencing was performed by the Genomic Sequencing and Analysis Facility at UT Austin, Center for Biomedical Research Support (RRID# SCR_021713). Computational analyses were performed using the Biomedical Research Computing Facility at UT Austin, Center for Biomedical Research Support (RRID#: SCR_021979). This work was supported by a US Department of Justice graduate fellowship, a UT Austin Graduate School Summer Fellowship, an IB graduate research grant, and Blair Scholarship fund in Zoology to IMC and NSF-IOS grant 1354942 and the William H. and Gladys G. Reeder Fellowship to HAH. IMC currently funded by Common Themes in Reproductive Diversity NIH 2T32HD049336.

