## Supplemental for "Preoptic transcriptomes during courtship reveal the molecular basis of behavioral coordination in a highly social cichlid"

AFFILITATIONS

[1]Department of Integrative Biology, The University of Texas at Austin, Austin, TX
[2]Institute for Neuroscience, The University of Texas at Austin, Austin, TX
[†]Current address: Department of Biology, Indiana University, Bloomington, IN

This file includes:

Legends for Datasets S1 to S4
Figures S1 to S6

**Dataset Legends**

**Dataset S1 (separate file)** Behavior and physiology sample data.

**Dataset S2 (separate file)** Correlation statistics for all pairwise behavior and physiology.

**Dataset S3 (separate file)** Gene count matrix per sample.

**Dataset S4 (separate file)** Gene module GO enrichment results.

**Dataset S5 (separate file)** Gene module membership from WGCNA for a list of known immediate-early genes and candidate genes related to social behavior.

**Dataset S6 (separate file)** Transcriptomic similarity z-score and z-score rank for both PCA distance and module eigengene distance for each partner pair.

### Supplemental Figures

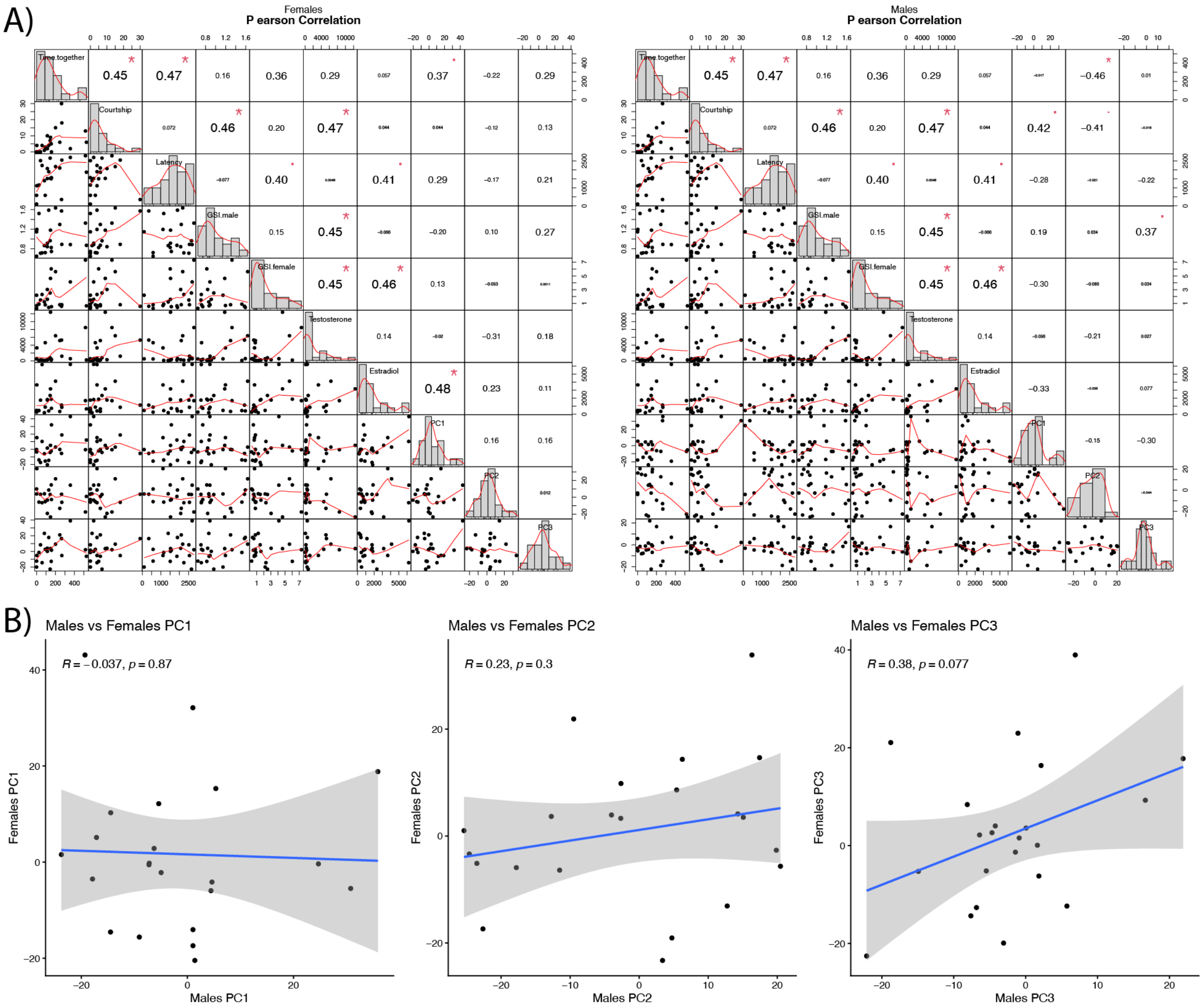


**Supplemental Figure 1.** Gene expression PCA. A) Correlation matrix of behavior and physiology traits for females (left) and males (right) with the first three PCs from the gene expression PCA. Variables include: Time spent together (Time.together), male courtship behavior (Courtship), latency for female to engage with male (Latency), male gonad size relative to body (GSI.male), female gonad size relative to body (GSI.female), male testosterone level (Testosterone), female estradiol level (Estradiol), and values from gene expression PCA for PC1, PC2, and PC3. B) Linear regression of the first three PCs of the gene expression PCA between partners.

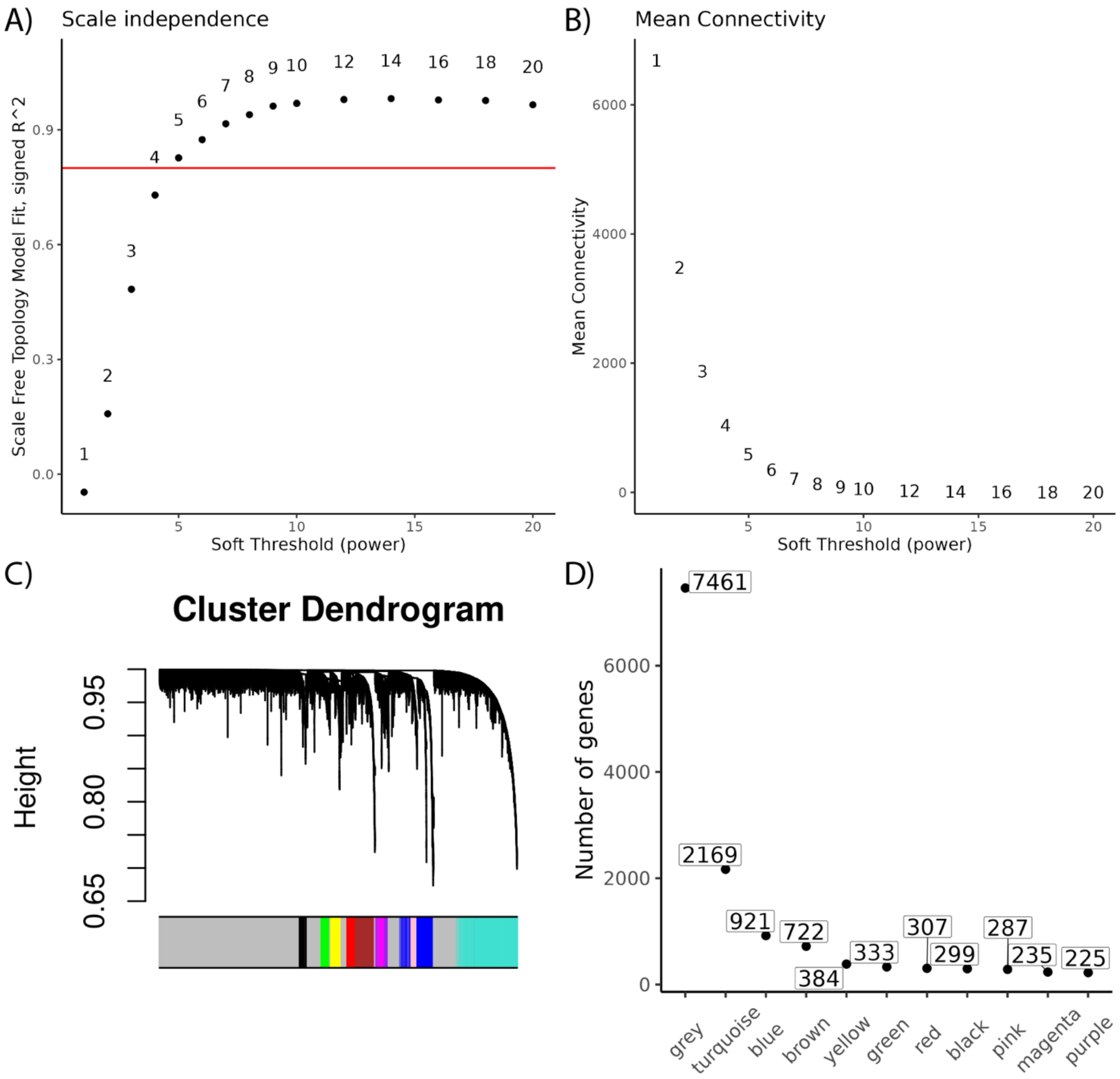


**Supplemental Figure 2.** WGCNA. A) Scale independence threshold for WGCNA. B) Mean connectivity threshold for WGCNA. C) Dendrogram of WGCNA resulting in ten gene modules. D) Number of genes contained in each module from WGCNA.

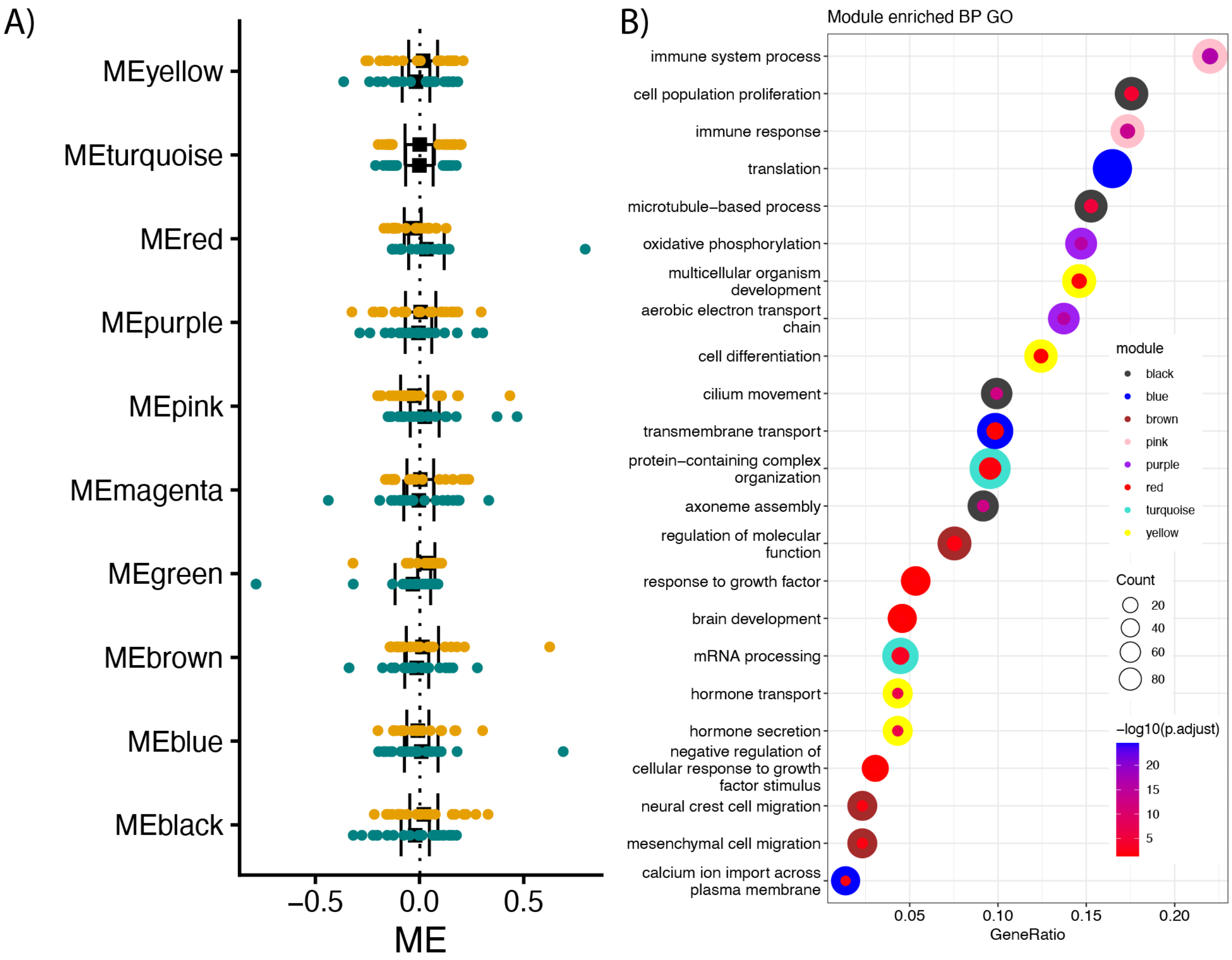


**Supplemental Figure 3.** A) Forest plot comparing module eigengene values of males (gold) and females (teal) for each gene module with mean and 95% confidence interval. B) GO enrichment analysis to identify significantly (FDR corrected p-value < 0.05) enriched GO terms for the GO category biological processes in each of the gene modules. Dotplot indicate top GO terms (y-axis), GO term gene ratio (x-axis), number of genes for that GO term (inside dot size), the FDR corrected p-value (inside dot color), and the module being tested (outside ring color). For each module, GO terms were selected to include the GO terms with the highest gene counts and most significant by adjusted p-value.

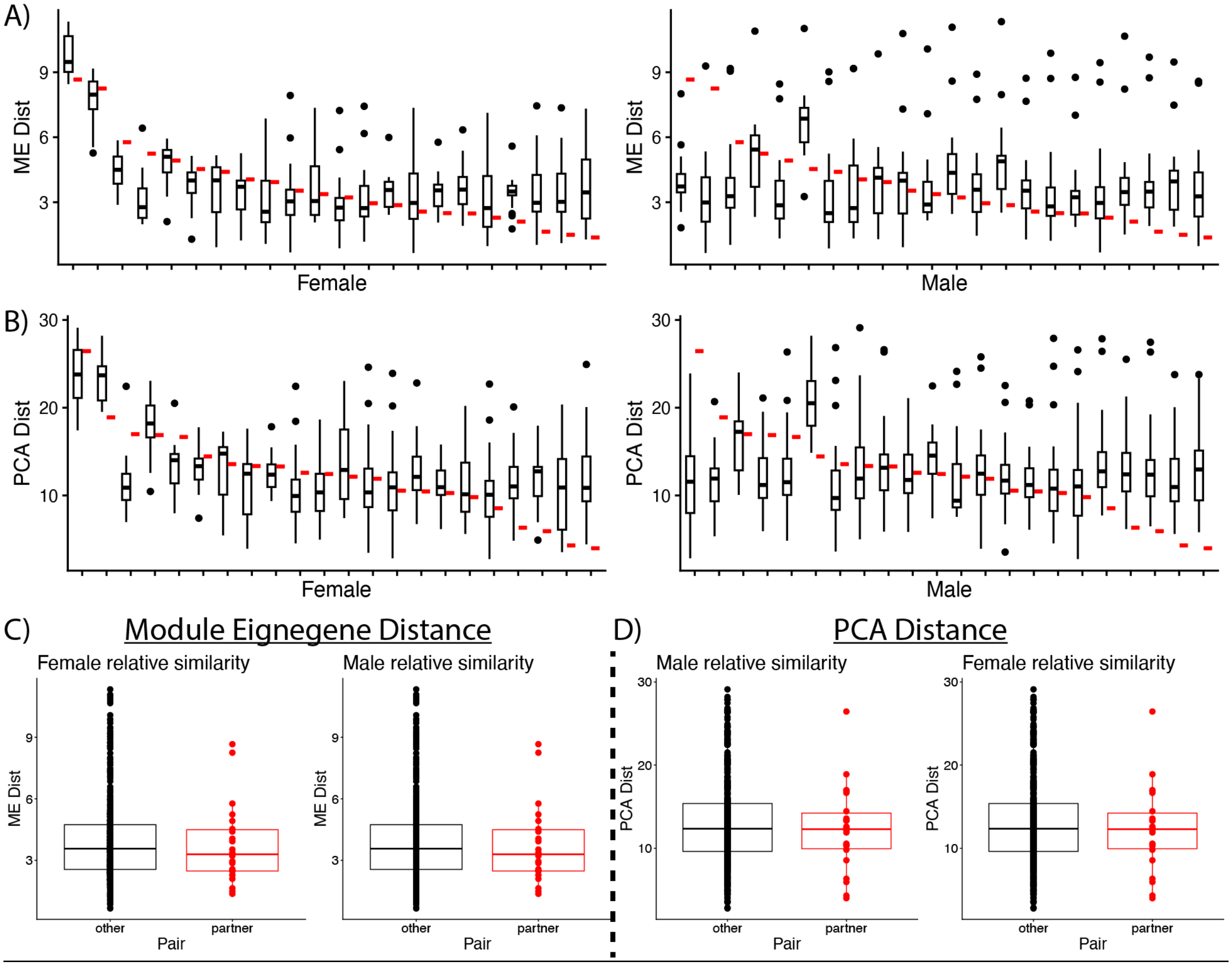


**Supplemental Figure 4.** A-B) Distribution of distance score (y-axis) for A) module eigengene distance and B) PCA distance for each individual (x-axis), for each female compared to every male (left) and each male compared to every female (right). The score of their partner is indicated in red. C-D) The total distribution of C) module eigengene distance and D) PCA distance for females (left) and males (right) between all other opposite-sex individuals (black) and their partner (red).

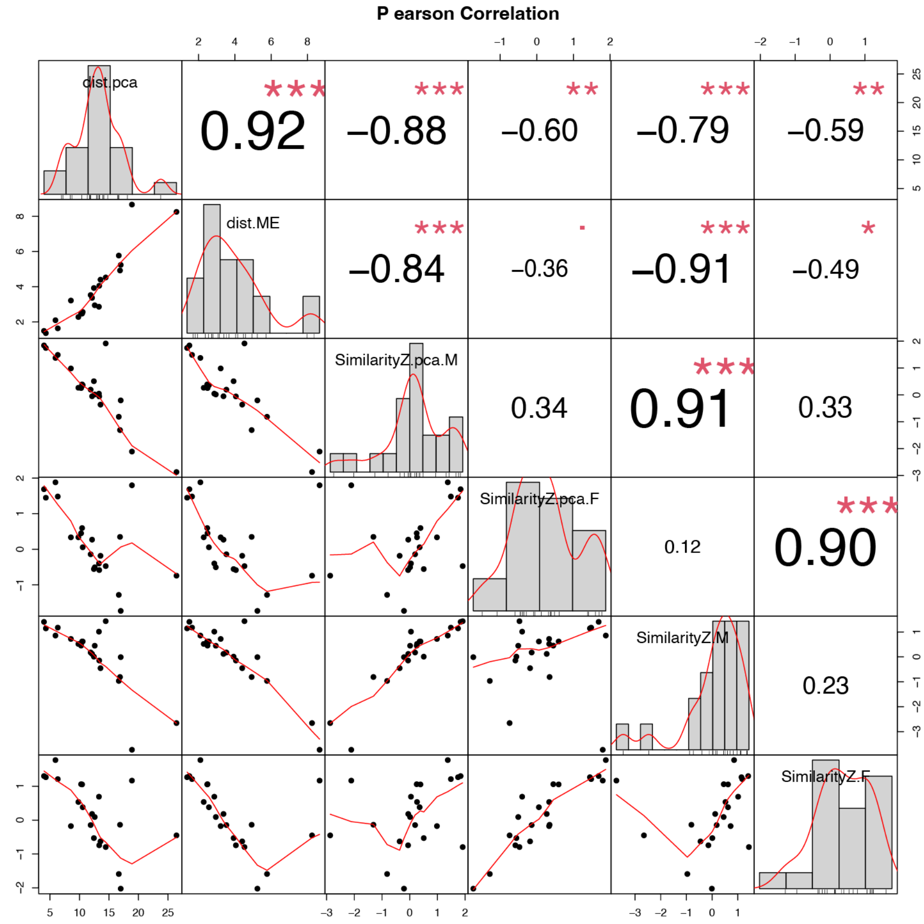


**Supplemental Figure 5.** Correlation of different transcriptomic similarity measures: gene expression weighted-Euclidean PCA distance (dist.pca), module eigengene distance (dist.ME), male PCA distance z-score (Similarity.Z.pca.M), female PCA distance z-score (Similarity.Z.pca.F), male module eigengene distance z-score (Similarity.Z.M), and female eigengene distance z-score (Similarity.Z.F).

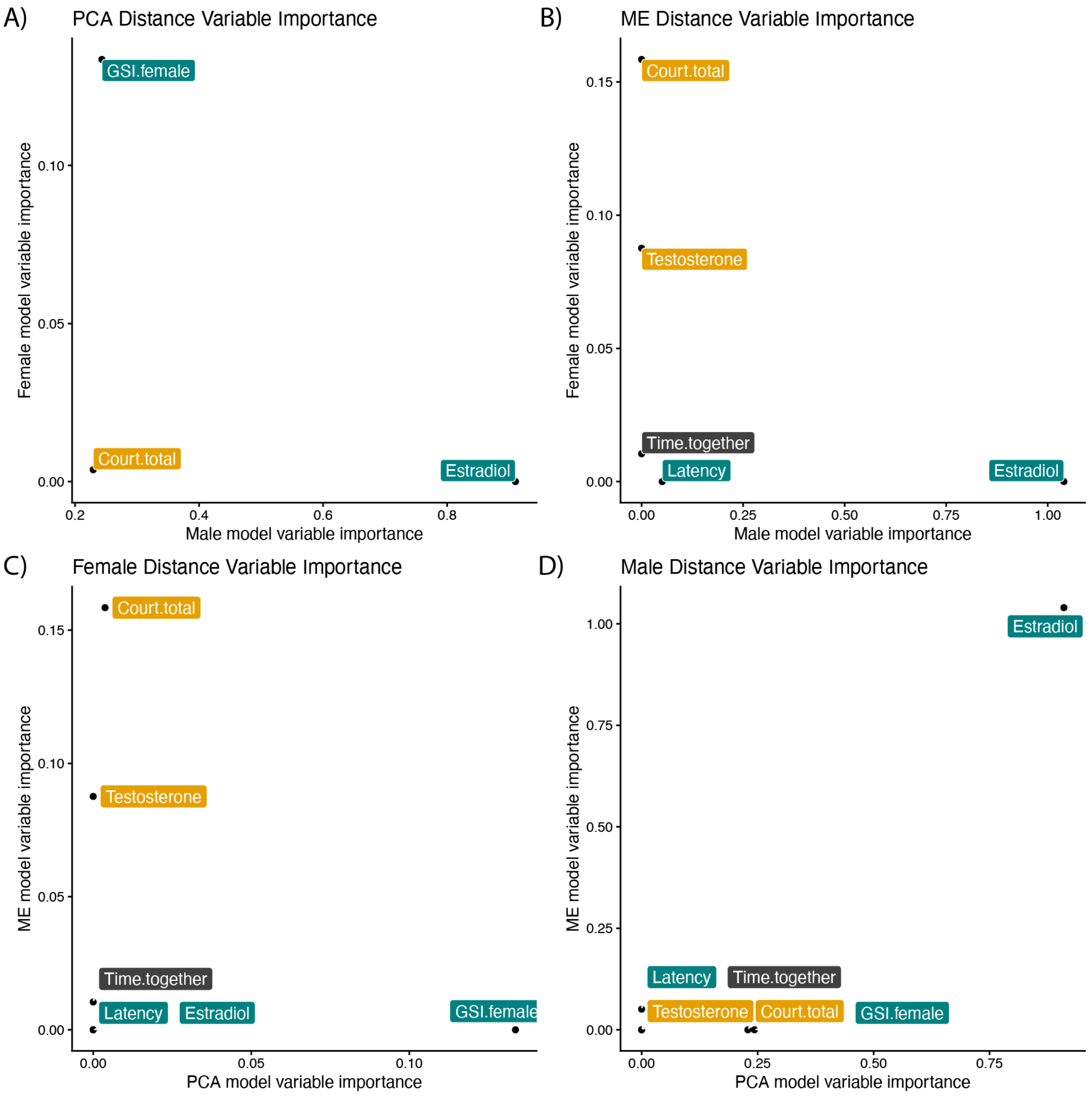


**Supplemental Figure 6.** A-B) Comparison of variable importance for male (x-axis) and female (y-axis) conditional inference tree models for A) PCA distance similarity and B) Module eigengene distance similarity. C-D) Comparison of variable importance for C) female and D) male (y-axis) conditional inference tree models for PCA distance similarity (x-axis) and Module eigengene distance similarity (y-axis).
